# High throughput screening of Leaf Economics traits in six wine grape varieties

**DOI:** 10.1101/2023.12.21.572933

**Authors:** Boya Cui, Rachel Mariani, Kimberley A. Cathline, Gavin Robertson, Adam R. Martin

## Abstract

Reflectance spectroscopy has become a powerful tool for non-destructive and high-throughput phenotyping in crops. Emerging evidence indicates that this technique allows for estimation of multiple leaf traits across large numbers of samples, while alleviating the constraints associated with traditional field- or lab-based approaches. While the ability of reflectance spectroscopy to predict leaf traits across species and ecosystems has received considerable attention, whether or not this technique can be applied to quantify within species trait variation have not been extensively explored. Employing reflectance spectroscopy to quantify intraspecific variation in functional traits is especially appealing in the field of agroecology, where it may present an approach for better understanding crop performance, fitness, and trait-based responses to managed and unmanaged environmental conditions. We tested if reflectance spectroscopy coupled with Partial Least Square Regression (PLSR) predicts rates of photosynthetic carbon assimilation (*A*_max_), Rubisco carboxylation (*V*_cmax_), electron transport (*J*_max_), leaf mass per area (LMA), and leaf nitrogen (N), across six wine grape (*Vitis vinifera*) varieties (Cabernet Franc, Cabernet Sauvignon, Merlot, Pinot Noir, Viognier, Sauvignon Blanc). Our PLSR models showed strong capability in predicting intraspecific trait variation, explaining 55%, 58%, 62%, and 64% of the variation in observed *J*_max_, *V*_cmax_, leaf N, and LMA values, respectively. However, predictions of *A*_max_ were less strong, with reflectance spectra explaining only 29% of the variation in this trait. Our results indicate that trait variation within species and crops is less well-predicted by reflectance spectroscopy, than trait variation that exists among species. However, our results indicate that reflectance spectroscopy still presents a viable technique for quantifying trait variation and plant responses to environmental change in agroecosystems.

## Introduction

Plant functional traits refer to the morphological, physiological, or phenological characteristics of plants that are readily measurable at an organismal scale, and influence the performance and response of individuals to environmental changes [1–4]. A considerable amount of effort has been directed towards understanding the extent, causes, and consequences of trait variation among plant species [5–10]. This body of literature has led to a deeper understanding of the key dimensions of functional trait variation that exist among the world’s plant species [6, 11].

Among the most well-studied dimensions of trait variation employed to describe and predict plant performance across resource availability gradients, is the “Leaf Economics Spectrum” (LES) [7–9]. The LES is a suite of six core leaf traits that covary among plant species including maximum photosynthetic assimilation (*A*_max_), leaf dark respiration rate (*R*_d_), leaf nitrogen (N) and phosphorus (P) concentrations, leaf mass per area (LMA), and leaf lifespan (LL). Taken together, LES trait expression defines how species vary across a continuum of life-history strategies, from fast-growing species characterized by rapid return on biomass investment, low structural investment, high leaf nutrient concentrations, and relatively short lifespans on one end, to resource-conserving species expressing the opposite suite of traits and by extension can be more resilient to resource limitation. Variation in LES traits largely owes to evolved trade-offs related to leaf biomechanics [12, 13], as well as evolved or plastic variation in physiological and leaf structural traits including stomatal and mesophyll conductance (*g*_s_ and *g*_m_, respectively), which in turn influence rates of maximum Rubisco carboxylation (*V*_cmax_), the electron transport (*J*_max_) [14–16].

Although much of the seminal work on trait variation is been based on interspecific comparisons, more recent research has focused on quantifying the extent and ecological implications of intraspecific trait variation [17–21]. Given the role that phenotypic plasticity and inheritable genetic variation play in governing plant ecophysiology and morphology, plant species can exhibit a high degree of intraspecific variation across a range of traits [21] and trait dimensions [17, 22]. Quantifying intraspecific trait variation is especially critical in agroecosystems where a relatively small number of plant species drive rates of ecosystem functioning on account of high abundances [23, 24]. Indeed, considerable interest and efforts have been dedicated to quantifying the causes and consequences of intraspecific variation in the traits that are directly responsible for crop growth, survival, and reproduction.

Though efforts to comprehensively assess intraspecific trait variation in a given plant species, especially crops, are often limited at the data collection phase of scientific enquiry. Traditionally, functional trait data are collected or derived from a combination of field and laboratory measurements, most of which can be laborious and time-consuming. This is especially true for “hard” traits [sensu 5] that are part of the LES such as *A*_max_ and *R*_d_ which are generated through point sampling of photosynthesis using portable infrared gas analyzers. Furthermore, traits that contribute to the physiological basis of LES trait variation, namely *V*_cmax_ and *J*_max_, rely on the execution and analysis of time-consuming photosynthetic CO_2_ response curves (*A*-*C*_i_) [reviewed by 25]. These methodological limitations to trait collection have at least in part motivated extensive research that evaluates how more easily-measured “soft” traits such as LMA can be used to predict “hard” physiological traits [5], especially in the context of Earth System Model parameterization [26, 27].

Reflectance spectroscopy has emerged as a central component of high-throughput phenotype assessments and related collection of physiological, chemical, and morphological trait data [28]. While multi- and hyperspectral sensors form a key component of remotely-sensed spectral diversity assessments at ecosystem scales [29–32], field-based reflectance spectroscopy offers an opportunity to rapidly amass species- or genotype-scale data on leaf physiological, chemical, and morphological traits including those forming the LES [33–35]. Specifically, using Partial Least Square Regression (PLSR) models [36], studies have reported strong predictive relationships between reflectance spectra and LES traits including *A*_max_, leaf N, LMA, and related physiological parameters including *V*_cmax_ and *J*_max_ [33, 37–39].

Spectroscopy coupled with PLSR models has been successful in estimating plant traits, particularly when using multi-species datasets that present a wide range of trait values and spectral profiles [33, 38, 40]. More recently, studies have begun employing these techniques to quantify and predict finer-scale intraspecific trait variation [41], including trait variation across individuals or genotypes of the same crop species [42–47]. Analyses on intraspecific trait variation—where trait values and spectra are more constrained—are less common vs. studies analyzing trait values and spectral signatures from a number of species differing in life-history strategies [33, 38] or agronomic profiles [40]. Furthermore, studies using reflectance spectroscopy to detect intraspecific trait variation in crops, commonly screen plants from a range of managed environmental conditions which further contributes a wider range of trait values [43]. While these results are promising, there remains uncertainty regarding whether or not these techniques are able to differentiate LES traits across individuals or genotypes of the same species, in agroecosystems where environmental conditions are more homogeneous.

Our study aims to contribute to the literature on high-throughput assessments of intraspecific trait variation, by evaluating the potential of reflectance spectroscopy to predict LES trait variations across multiple wine grape (*Vitis vinifera*) varieties: one of the most common crops that holds substantial agricultural and economic values. In this study, we hope to determine whether PLSR models can reliably estimate photosynthetically important functional traits in wine grapes from reflectance spectroscopy data across six cultivated varieties.

## Materials and Methods

### Study site

We collected LES and related trait and spectral reflectance data for six of the most common wine grape varieties—Cabernet Franc, Cabernet Sauvignon, Merlot, Pinot Noir, Sauvignon Blanc, Viognier—at the Niagara College Teaching Vineyard, Niagara-on-the-Lake, Ontario. The site is an operational vineyard characterized as non-irrigated, with imperfectly drained silty clays overlaying clay loam till mixed with poorly drained lacustrine heavy clay, and uniformly tilled and sprayed [48, 49]. All trait and reflectance data were collected during the fruit setting stage (at our site, from June 6-17, 2022) between 6:00-12:00. For each variety, we sampled 30 vines evenly distributed across three planting rows, which were roughly 10 meters apart from each other within one row, totalling *n*=180 individual vines. One leaf on each vine was selected from the uppermost segment of the individual for data collection, with all leaves being fully exposed, newly developed, fully expanded, and free of any damage [50].

### Functional trait data collection

Trait data in our study included *A*_max_, *V*_cmax_, and *J*_max_, leaf N concentrations, and LMA. First, *V*_cmax_, *J*_max_, and *A*_max_ data were collected in the field using a LI-6800 Portable Photosynthesis System (Licor Bioscience, Lincoln, Nebraska, USA). We first performed an *A-C_i_*curve on each leaf using the Dynamic Assimilation Technique (DAT) [25, 51, 52] in order to estimate rates of *V*_cmax_ and *J*_max_. For each curve, CO_2_ assimilation rates on a per leaf area basis (*A*_area_; *μ*mol CO_2_ m^-2^ s^-1^) were logged every 4 seconds across continuously ramping CO_2_ concentrations, with a ramp rate of 100 *μ*mol mol^-1^ min^-1^ [consistent with recommendations by 52, 53] beginning at 5 *μ*mol mol^-1^ CO_2_ and concluding at 1700 *μ*mol mol^-1^ CO_2_. Otherwise, conditions in the leaf chamber were set to a photosynthetic photon flux density (PPDF) of 1500 *μ*mol m^-2^ s^-1^ of photosynthetically active radiation (PAR; 400-700 nm), 50% relative humidity, leaf vapour pressure deficits of 1.7 KPa, and leaf temperatures of 25 °C. Furthermore, CO_2_ and H_2_O sensors were readjusted using the range match function after every five leaf measurements, and each DAT *A-C_i_* curve required approximately 10 minutes, including a 60-120 second acclimation period [25]. Following the completion of each *A*-*C*_i_ curve, we then allowed leaves to acclimate to ambient conditions for ∼10 minutes. Then, we collected steady-state *A*_max_ values for each leaf at the same environmental conditions as mentioned above with a constant CO_2_ concentration at 420 ppm. We logged steady-state gas *A*_max_ values after leaves were allowed to stabilize for 5-10 minutes.

Immediately following gas exchange measurements, we used an HR1024i full spectrum portal field spectroradiometer (Spectra Vista Corporation, Poughkeepsie NY, USA) to collect reflectance spectra for each leaf. This instrument is a full-range spectroradiometer (350-2500 nm) with a spectral resolution of ≤3.5 nm (350-1000 nm), ≤9.5 nm (1000-1800 nm), and ≤ 6.5 nm (1800-2500 nm), outfitted with an LC-RP Pro leaf clip that includes a calibrated internal light source. Reflectance spectra were collected at the same location on the adaxial side of each leaf from which *A*-*C*_i_ and steady state gas exchange were performed, with integration times set to 2 seconds, and reference spectra taken on a white Spectralon standard prior to each measurement.

Once physiological and reflectance data were acquired, we collected and transported individual leaves to the University of Toronto Scarborough for quantification of LMA and leaf N concentrations. First, we removed all petioles, and the fresh area of all leaves was quantified using an LI-3100C leaf area meter (Licor Bioscience, Lincoln, Nebraska, USA), and then dried for 48 hours to constant mass. Dried leaves were then weighed and LMA was calculated as mass/ area. Finally, dried leaves were ground to a fine and homogeneous powder using a MM400 Retsch ball mill (Retsch Ltd., Hann, Germany), and a LECO CN 628 elemental analyzer (LECO Instruments, Ontario, Canada) was used to determine leaf N concentrations on ∼0.1 grams of powdered tissue.

### Data analysis

R Statistical Software v. 4.2.0 (R Foundations for Statistical Computing, Vienna, Austria) was used for all data analysis. First, we fit the Farquhar, von Caemmerer and Berry (FvCB) model to each individual *A*-*C*_i_ curve, using the ‘fitaci’ function in the ‘plantecophys’ R package [54], in order to estimate rates of *V*_cmax_ and *J*_max_, along with their standard errors. In this procedure, these models were fit using non-linear least square regression [54], such that *V*_cmax_ and *J*_max_ were corrected to 25 °C, and *V*_cmax_ and *J*_max_ are considered apparent as mesophyll conductance was assumed to be infinite. These data were merged with other traits, and the distribution of each individual trait was assessed using the ‘fitdist’ function in the *‘*fitdistrplus*’* R package [55]. Traits were determined to be either normally or log-normally distributed (as per the highest log-likelihood value) and transformed data was employed in further analyses in accordance with these results. We then performed an analysis of variance (ANOVA) to test for significant trait differences across varieties.

We then followed the methods described by Burnett et al. [36] to evaluate how reflectance spectra predicted trait values across our dataset, using a PLSR modelling approach. All PLSR models included reflectance spectral data from the 500-2400 nm wavelength range, and aimed to predict either non-transformed or log-transformed trait data, as informed by our distribution fitting procedure. For each PLSR model, the spectra-trait dataset was split into a calibration dataset (which included 80% of all data points) and a validation dataset (comprised of the remaining 20% of data). Since we were explicitly interested in testing the ability of reflectance spectra to quantity variation in leaf traits across grapes broadly, and the ability to differentiate varieties, we performed and analyzed two data splits. First, datasets were split into calibration vs. validation according to variety identity, such that both the validation and calibration datasets had approximately equal proportions of trait and spectra data from all varieties. Second, we used a completely randomized data split, whereby the proportion of data across varieties was allowed to vary randomly.

Using the calibration datasets, we then used the ‘find_optimal_components’ function in the ‘spectratrait’ R package [56] to determine the optimal number of components used in the final PLSR model, based on the minimization of the prediction residual sum of squares (PRESS) statistic. For each trait, a PLSR model was fitted from the calibration dataset using the leave-one-out cross-validation (LOO) procedure, specified with the ‘plsr’ function in the ‘pls’ R package [57]. Model performance was then assessed with the validation datasets as an external validation, in which the predicted values and the observed values in the validation dataset were compared.

For the final models, we used the validation coefficient of determination (*r*^2^), root mean squared error of prediction (RMSE), and percent root mean squared error of prediction (%RMSE) as metrics to illustrate model fits.

To further evaluate the model performance, we used the model coefficients and variable influences on projection (VIP) values to explore the effect of different areas of the spectra on predicting the trait variable. Following this, we performed a jackknife permutation analysis to assess model uncertainty, using the jackknife argument of the ‘plsr’ function in the ‘pls’ R package [57]. The resulting jackknife coefficients were then compared to that of the full model. And finally, using the full model and jackknife permutation outputs, the mean, and 95% confidence and prediction intervals were calculated for each predicted trait value from the validation dataset.

## Results

### Reflectance spectroscopy for predicting within-variety leaf traits

Leaf traits measured here all varied significantly as a function of variety identity (*p*<0.001 in all cases). Specifically, across the entire dataset, physiological traits were most variable, with *A*_max_ ranging from 3.8-29.0 *μ*mol CO_2_ m^-2^ s^-1^ (CV=34.8), *V*_cmax_ from 28.9-131.7 *μ*mol m^-2^ s^-1^ (CV=27.5), and *J*_max_ from 60.3-253.1 *μ*mol m^-2^ s^-1^ (CV=25.8). In comparison, LMA and leaf N also varied significantly across varieties, though these traits were less variable with LMA ranging from 52.8-101.8 g m^-2^ (CV=12.8) and leaf N from 2.04-4.39% (CV=13.7). All reflectance spectra presented generally the same shape, with a few Cab. Franc individuals situated closer to the lower range, Merlot and Pinot Noir closer to the upper range, and others in and around the 95% confidence interval (Figure 1).

**Figure 1.**
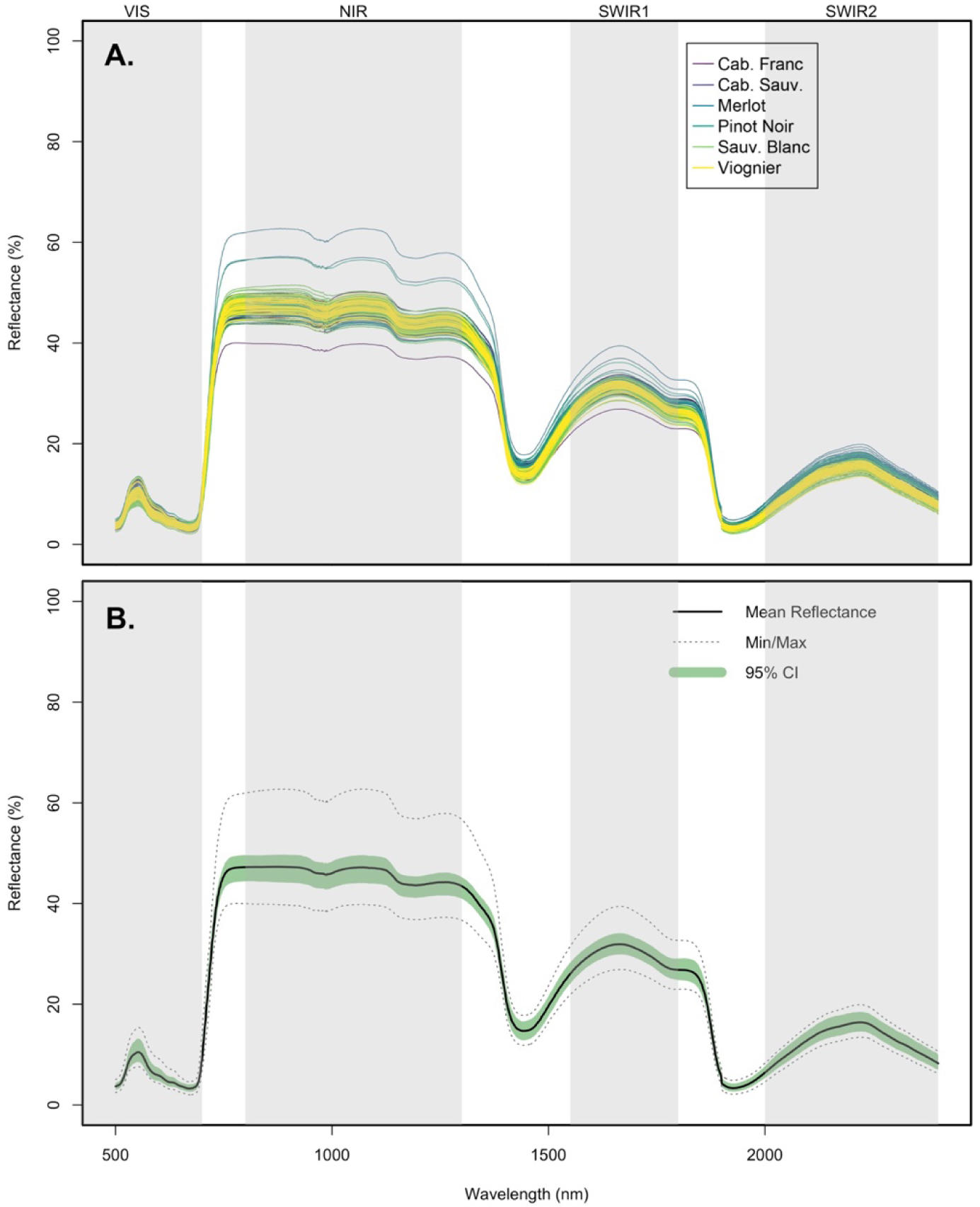
Reflectance spectra of 179 wine grape leaves plotted A) individually with six wine grape varieties specified, and B) all together with mean, range, and 95% confidence interval estimates. All spectral data were trimmed to the 500-2400 nm range where the PLSR models were built from. The grey shaded areas indicate different spectral regions: Visible Spectrum (VIS), Near Infrared (NIR), Short Wave Infrared 1 (SWIR1), and Short Wave Infrared 2 (SWIR2).

When calibration vs. validation data were evenly split across varieties (i.e., 80% of each variety allocated to each dataset), reflectance spectra and PLSR models explained between 18- 64% of the variation in wine grape traits (Table 1, Figure 2). Specifically, physiological traits including *A*_max_, *J*_max_, and *V*_cmax_ were predicted by 4-5 spectral components which cumulatively explained 18%, 44%, and 30% of the variation in these traits, respectively. In these cases, model %RMSE values ranged from 21.6% in *A*_max_ models, 24.1% in *V*_cmax_ models, and 18.9% in *J*_max_ models. Comparatively, reflectance spectra and PLSR models expressed stronger predictive ability towards log-LMA and leaf N, with models (r2) explaining 64% (%RMSE=14.3) and 62% (%RMSE=15.2%) of the variation, respectively (Table 1, Figure 2).

**Figure 2.**
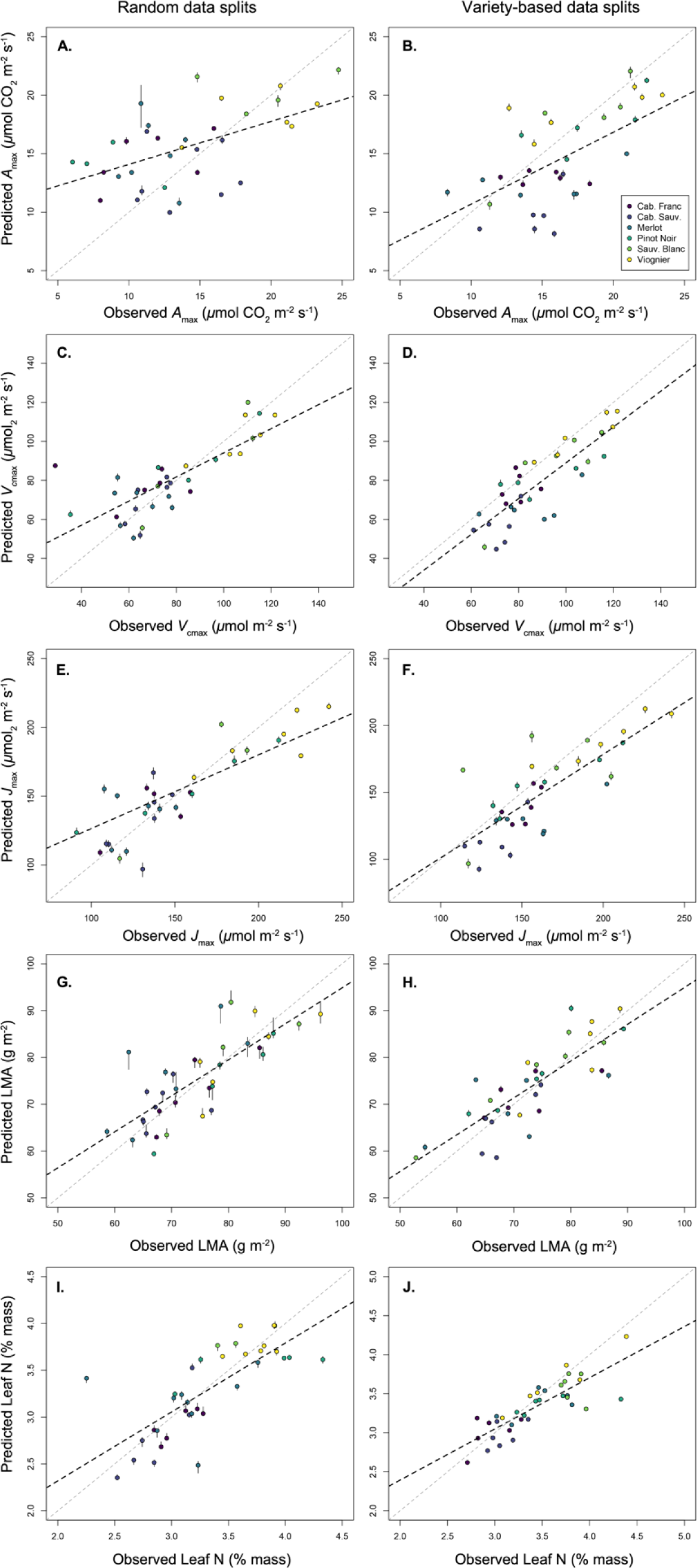
Results of partial least squares regression (PLSR) models predicting leaf physiological, chemical, and morphological traits in six wine grape varieties. Shown here are the data points used to validate the models (*n*=36 in all cases) fitted to a set of calibration data points (*n*=176- 178; see Table 1). Calibration and validation datasets were selected on the basis of a fully randomized data split (left panels), and a data split where all six varieties were equally represented in the calibration datasets (right panels). Dashed black lines represent linear model fits between observed vs. expected trait values, while the dotted gray lines represent a 1:1 relationship.

**Table 1.** Partial least squares regression model fits evaluating the ability of reflectance spectra to explain variation in leaf traits measured on six wine grape varieties. Presented here are results from two different modelling approaches which divide our total sample into calibration (80% of our data) vs. validation (20% of our data) datasets. In the results associated with the “Variety” approach, calibration and validation data both included approximately the same proportions of observations from all varieties, while the “Random” approach made this division randomly. Here, *n*_obs_ refers to the total observations in our dataset for a given trait, which entails a correspond sample size in the validation dataset (*n*_val_). For each model we present the number of components derived from reflectance spectra that were included in the final predictive model (n_comp_), along with the root mean square error (RSME), *r*^2^ value, and %RMSE for the final predictive model. All models were based on Trait acronyms are as follows: light saturated photosynthetic rate (*A*_max_), maximum velocity of Ribulose 1,5-bisphosphate (RuBP) carboxylation (*V*_cmax_), maximum rate of electron transport (*J*_max_), leaf mass per area (LMA), leaf nitrogen (N) concentration.

| Data split | Trait | $n_{\text{obs}}$ | $n_{\text{val}}$ | $n_{\text{comp}}$ | RMSE | $r^2$ | %RMSE |
| --- | --- | --- | --- | --- | --- | --- | --- |
| Variety | $A_{\text{max}}$ | 178 | 36 | 4 | 3.9 | 0.18 | 21.63 |
| | $V_{\text{cmax}}$ | 177 | 36 | 5 | 14.62 | 0.3 | 24.1 |
| | $J_{\text{max}}$ | 177 | 36 | 4 | 24.22 | 0.44 | 18.88 |
|  | log-LMA | 178 | 36 | 10 | 0.08 | 0.64 | 14.27 |
|  | Leaf N | 176 | 36 | 9 | 0.25 | 0.62 | 15.16 |
| Random | $A_{\text{max}}$ | 178 | 36 | 1 | 4.42 | 0.29 | 19.25 |
| | $V_{\text{cmax}}$ | 177 | 36 | 9 | 14.47 | 0.58 | 15.6 |
| | $J_{\text{max}}$ | 177 | 36 | 8 | 28.35 | 0.55 | 15.6 |
|  | log-LMA | 178 | 36 | 13 | 0.08 | 0.53 | 16.44 |
|  | Leaf N | 176 | 36 | 11 | 0.33 | 0.5 | 15.92 |

The predictive power of PLSR models was sensitive to the configuration of calibration and validation datasets, though general trends were nuanced. When calibration and validation datasets were comprised of varieties in random proportions, physiological traits were better predicted than in datasets where variety proportions were equal. Specifically, in randomized data splits, *A*_max_ model *r*^2^=0.29, *V*_cmax_ *r*^2^=0.58, and *J*_max_ *r*^2^=0.55, all of which were higher vs. the same models in variety-weighted data splits. Alternatively, PLSR models for log-LMA and leaf N had lower predictive power when calibration and validation datasets were randomly created, with *r*^2^ values of 0.53 and 0.5, respectively (Table 1, Figure 2). In all cases, the number of spectral components retained in the final PLSR models also differed depending on the nature of calibration and validation dataset construction.

The impact of the data splitting method is also observed in the model regression coefficient trends, which reflect the contribution of certain wavelengths to trait prediction. For physiological traits, the shapes of regression coefficient trends are similar within the same splitting method, but distinctly different between splitting methods (Figure 3). Here we ignore the random split model of *A*_max_ from this comparison, due to its limited number of model components. On the other hand, splitting data randomly or proportionally across varieties did not influence the regression coefficient distributions of log-LMA or leaf N (Figure 3). VIP scores of the models suggest similar wavelength regions of importance for model prediction across different traits, regardless of data splitting methods (Figure 4).

**Figure 3.**
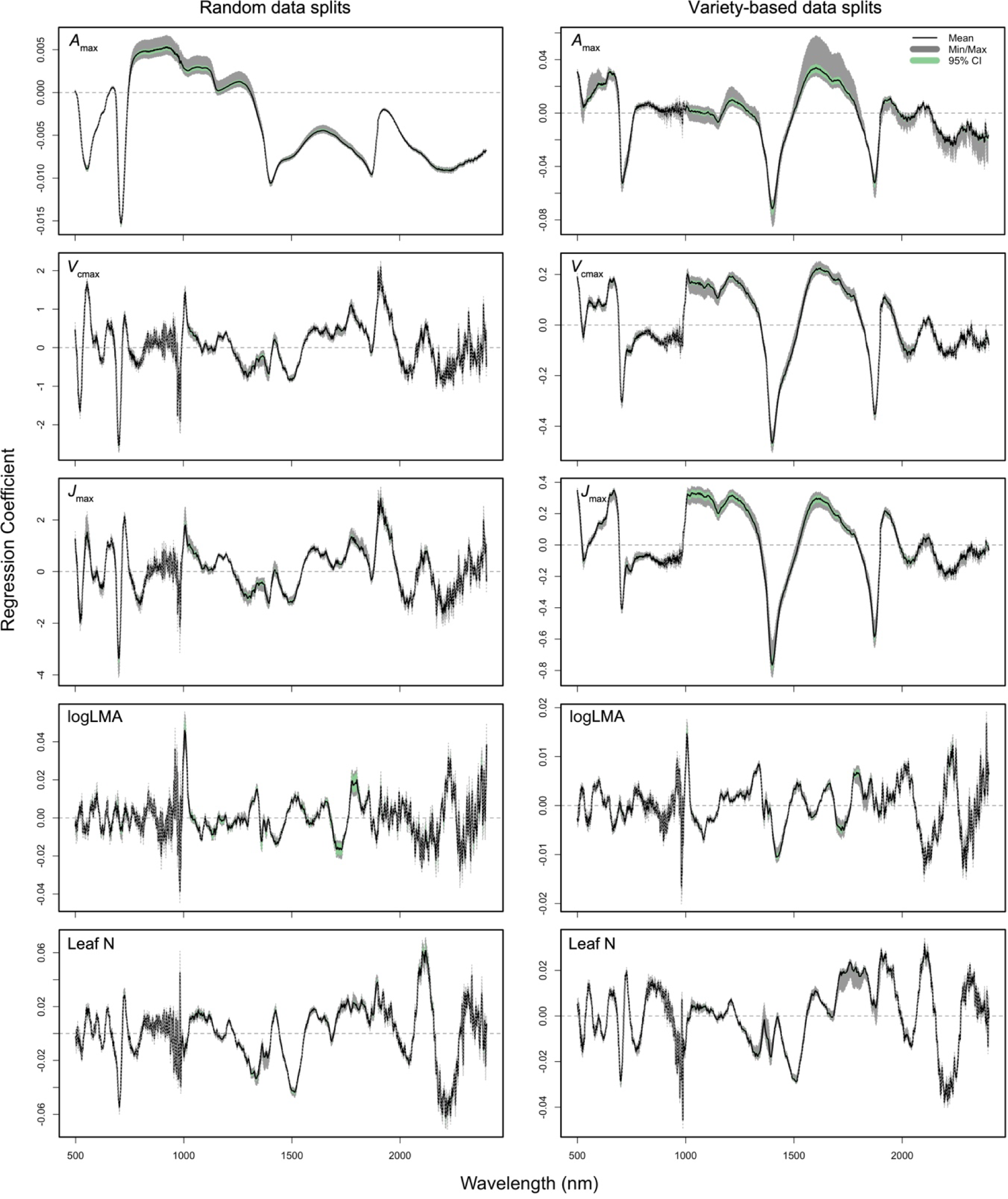
Jackknife regression coefficients of the PLSR models of *A*_max_, *V*_cmax_, *J*_max_, log-LMA, and leaf N, based on the calibration data. The dashed horizontal line in each panel indicates where the coefficient is zero. The black curve represents the mean, the grey area represents the range, and the green area represents the 95% confidence interval.

**Figure 4.**
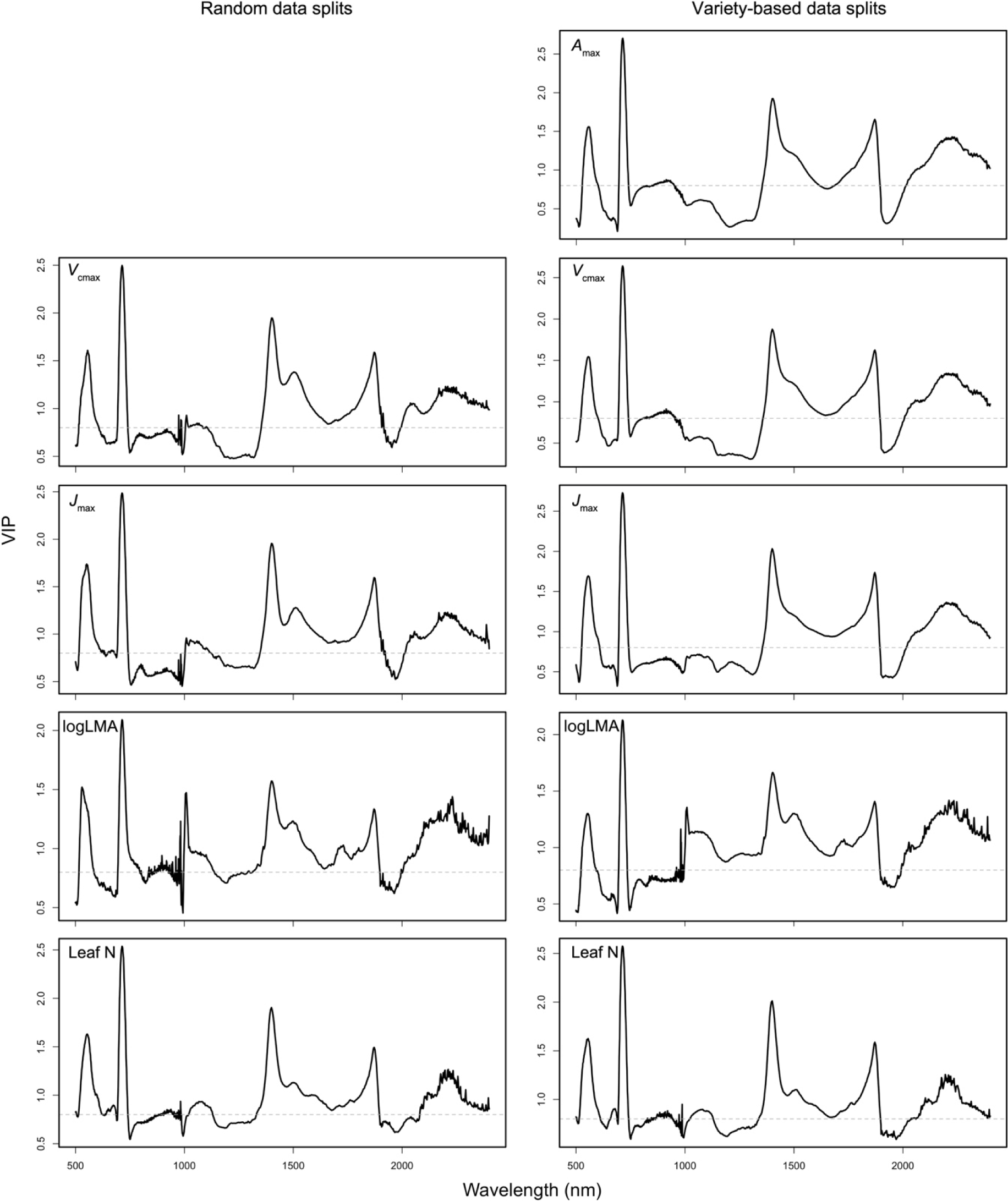
Variable influences on projection (VIP) scores of the final PLSR models of *A*_max_, *V*_cmax_, *J*_max_, log-LMA, and leaf N. The dashed horizontal line in each panel indicates where the VIP score is 0.8. The *A*_max_ model using random data split method had one component and therefore did not generate valid VIP scores.

## Discussion

Our findings contribute to the growing literature that reflectance spectroscopy is well-equipped to detect trait variation within and among plant species [27, 33, 35, 38, 40, 41, 45]. A considerable proportion of earlier work in this area focused on quantifying the interspecific trait variation that exists among plants of different functional types, that differ widely their evolutionary histories and trait diversity [e.g., 33, 35]. To this end, previous studies have indicated that reflectance spectroscopy is better equipped to explain trait variation, in situations where trait values within calibration and validation datasets vary more widely [38]. This tendency positions these techniques for rapid trait estimation in natural ecosystems [32], with many such studies reporting a high predictive ability of PLSR models in quantifying interspecific trait variation. Though a recent renewed focus on the importance of intraspecific trait variation in driving ecosystem functioning [20, 21], along with applications of these techniques in certain fields including agroecology, necessitates quantifying and disentangling the drivers of finer-scale trait variation that generally exists within species [24].

In this regard, our results show the strong predictive power of PLSR models to capture between 50-64% of the within-species trait variation in wine grapes, for key LES and related traits including *V*_cmax_, *J*_max_, log-LMA, and leaf N (Table 1, Figure 2). Previous studies that examined within-species trait variation using PLSR approaches have yielded broadly similar results. For example, Meacham-Hensold et al. [46] reported PLSR models that explained 60%, 59%, and 83% of the variation in *V*_cmax_, *J*_max_, and leaf N, respectively, across six tobacco (*Nicotiana tabacum*) genotypes, though when three additional genotypes and larger sample sizes were included in analyses, these PLSR model *r*^2^ values increased to 0.61 for *V*_cmax_, and 0.62 for *J*_max_ in the validation dataset. Similarly, Fu et al. [47] modelled photosynthetic traits of six tobacco genotypes using PLSR methods, and reported similar *r*^2^ values (0.60 and 0.56) for *V*_cmax_ and *J*_max_, respectively.

Other single-species studies that applied reflectance spectroscopy and PLSR models to predict leaf traits across experimental treatments or environmental gradients have also presented similar results. For example, Yendrek et al. [43] found reflectance spectra were strong predictors of leaf N (*r*^2^= 0.92-0.96) and *V*_cmax_ (*r*^2^=0.56-0.65) of maize (*Zea mays*) genotypes grown across gradients of ozone and soil N availability. Finally, in an analysis that screened over 200 genotypes of wheat (*Triticum aestivum*, *T. turgidum*, and triticale germplasm) from six sets of experiments, Silva-Perez et al. [42] included detected high predictive power of PLSR models, with *r*^2^ values ranging from 0.70-0.89 for leaf N, LMA, *V*_cmax_, and *J*_max_. Though in this same experiment, consistent with our results CO_2_ assimilation rates were relatively poorly captured by PLSR models: in our analysis, the *r*^2^ for models for *A*_max_ were 0.18-0.29, vs. *r*^2^ values of 0.49 in Silva-Perez et al. [42].

In addition to model diagnostics alone, in our analysis, PLSR models generally support the same inferences surrounding the comparative trait biology of wine grape varieties (relative to observed trait data). Specifically, our previous analysis of LES trait variation—with trait data observed in the field using traditional gas exchange and analytical chemistry techniques—found that white grape varieties Sauvignon Blanc and Viognier occupy the “resource-acquiring” end of an intraspecific LES in wine grapes (characterized by high rates of *A*_max_, *V*_cmax_, *J*_max_, leaf N, and low LMA), while red varieties (Cabernet Franc, Cabernet Sauvignon, Merlot) define the “resource-conserving” end of the wine grape LES (characterized by low *A*_max_, *V*_cmax_, *J*_max_, leaf N, and high LMA). Our PLSR models support this same general trend (Figure 1), with white varieties expressing predictions that indicate resource-acquiring trait values.

Our analysis contributes evidence that reflectance spectroscopy and PLSR modelling approaches, can be used to 1) directly predict intraspecific trait variation with a relatively high degree of accuracy, and 2) differentiate intraspecific variation in life-history strategies in plants. Though our analysis here is based on a small subset of the 1,000s of wine grape varieties that exist globally [58]. Therefore, expanding this work to include a greater number of wine grape genotypes and trait values [cf. 41, 42], presents a viable opportunity to more rapidly screen trait expression in one of the world’s most economically important crops.

## Acknowledgments

The authors acknowledge both Mitchell Madigan and Lauren Miller for their assistance with field data collection. This study was supported by a Discovery Grant to A.R.M. from the Natural Sciences and Engineering Research Council of Canada, and by the University of Toronto Scarborough’s (UTSC) Sustainable Food and Farming Futures (SF3) Cluster under UTSC’s Clusters of Scholarly Prominence Program.

## References

1. Díaz S, Cabido M. Plant functional types and ecosystem function in relation to global change. Journal of vegetation science. 1997;8(4):463–74.

2. Funk JL, Larson JE, Ames GM, Butterfield BJ, Cavender-Bares J, Firn J, et al. Revisiting the H oly G rail: using plant functional traits to understand ecological processes. Biological Reviews. 2017;92(2):1156–73.

3. Lavorel S, Garnier E. Predicting changes in community composition and ecosystem functioning from plant traits: revisiting the Holy Grail. Functional ecology. 2002;16(5):545–56.

4. Westoby M, Wright IJ. Land-plant ecology on the basis of functional traits. Trends in ecology & evolution. 2006;21(5):261–8.

5. Diaz S, Hodgson J, Thompson K, Cabido M, Cornelissen JH, Jalili A, et al. The plant traits that drive ecosystems: evidence from three continents. Journal of vegetation science. 2004;15(3):295–304.

6. Díaz S, Kattge J, Cornelissen JH, Wright IJ, Lavorel S, Dray S, et al. The global spectrum of plant form and function. Nature. 2016;529(7585):167-71.

7. Reich PB, Ellsworth DS, Walters MB, Vose JM, Gresham C, Volin JC, et al. Generality of leaf trait relationships: a test across six biomes. Ecology. 1999;80(6):1955–69.

8. Wright IJ, Reich PB, Cornelissen JH, Falster DS, Garnier E, Hikosaka K, et al. Assessing the generality of global leaf trait relationships. New phytologist. 2005;166(2):485–96.

9. Wright IJ, Reich PB, Westoby M, Ackerly DD, Baruch Z, Bongers F, et al. The worldwide leaf economics spectrum. Nature. 2004;428(6985):821-7.

10. Westoby M, Falster DS, Moles AT, Vesk PA, Wright IJ. Plant ecological strategies: some leading dimensions of variation between species. Annual review of ecology and systematics. 2002;33(1):125–59.

11. Carmona CP, Bueno CG, Toussaint A, Träger S, Díaz S, Moora M, et al. Fine-root traits in the global spectrum of plant form and function. Nature. 2021;597(7878):683-7.

12. Shipley B, Lechowicz MJ, Wright I, Reich PB. Fundamental trade-offs generating the worldwide leaf economics spectrum. Ecology. 2006;87(3):535–41.

13. Blonder B, Violle C, Bentley LP, Enquist BJ. Venation networks and the origin of the leaf economics spectrum. Ecology letters. 2011;14(2):91–100.

14. Xiong D, Flexas J. Leaf economics spectrum in rice: leaf anatomical, biochemical, and physiological trait trade-offs. Journal of Experimental Botany. 2018;69(22):5599–609.

15. Onoda Y, Wright IJ, Evans JR, Hikosaka K, Kitajima K, Niinemets Ü, et al. Physiological and structural tradeoffs underlying the leaf economics spectrum. New Phytologist. 2017;214(4):1447–63.

16. Onoda Y, Wright IJ. The leaf economics spectrum and its underlying physiological and anatomical principles. The leaf: a platform for performing photosynthesis. 2018:451–71.

17. Anderegg LD, Berner LT, Badgley G, Sethi ML, Law BE, HilleRisLambers J. Within-species patterns challenge our understanding of the leaf economics spectrum. Ecology letters. 2018;21(5):734–44.

18. Bolnick DI, Amarasekare P, Araújo MS, Bürger R, Levine JM, Novak M, et al. Why intraspecific trait variation matters in community ecology. Trends in ecology & evolution. 2011;26(4):183–92.

19. Violle C, Enquist BJ, McGill BJ, Jiang L, Albert CH, Hulshof C, et al. The return of the variance: intraspecific variability in community ecology. Trends in ecology & evolution. 2012;27(4):244–52.

20. Westerband A, Funk J, Barton K. Intraspecific trait variation in plants: a renewed focus on its role in ecological processes. Annals of botany. 2021;127(4):397–410.

21. Siefert A, Violle C, Chalmandrier L, Albert CH, Taudiere A, Fajardo A, et al. A global meta-analysis of the relative extent of intraspecific trait variation in plant communities. Ecology letters. 2015;18(12):1406–19.

22. Isaac ME, Martin AR, de Melo Virginio Filho E, Rapidel B, Roupsard O, Van den Meersche K. Intraspecific trait variation and coordination: Root and leaf economics spectra in coffee across environmental gradients. Frontiers in plant science. 2017;8:1196.

23. Milla R, Osborne CP, Turcotte MM, Violle C. Plant domestication through an ecological lens. Trends in ecology & evolution. 2015;30(8):463–9.

24. Martin AR, Isaac ME. Plant functional traits in agroecosystems: a blueprint for research. Journal of Applied Ecology. 2015;52(6):1425–35.

25. Saathoff AJ, Welles J. Gas exchange measurements in the unsteady state. Plant, Cell & Environment. 2021;44(11):3509–23.

26. Reich P, Ellsworth D, Walters M. Leaf structure (specific leaf area) modulates photosynthesis–nitrogen relations: evidence from within and across species and functional groups. Functional Ecology. 1998;12(6):948–58.

27. Serbin SP, Wu J, Ely KS, Kruger EL, Townsend PA, Meng R, et al. From the Arctic to the tropics: multibiome prediction of leaf mass per area using leaf reflectance. New Phytologist. 2019;224(4):1557–68.

28. Fahlgren N, Gehan MA, Baxter I. Lights, camera, action: high-throughput plant phenotyping is ready for a close-up. Current opinion in plant biology. 2015;24:93–9.

29. Asner GP, Martin RE, Anderson CB, Knapp DE. Quantifying forest canopy traits: Imaging spectroscopy versus field survey. Remote Sensing of Environment. 2015;158:15–27.

30. Singh A, Serbin SP, McNeil BE, Kingdon CC, Townsend PA. Imaging spectroscopy algorithms for mapping canopy foliar chemical and morphological traits and their uncertainties. Ecological Applications. 2015;25(8):2180–97.

31. Chadwick KD, Brodrick PG, Grant K, Goulden T, Henderson A, Falco N, et al. Integrating airborne remote sensing and field campaigns for ecology and Earth system science. Methods in Ecology and Evolution. 2020;11(11):1492–508.

32. Asner GP, Knapp DE, Anderson CB, Martin RE, Vaughn N. Large-scale climatic and geophysical controls on the leaf economics spectrum. Proceedings of the National Academy of Sciences. 2016;113(28):E4043–E51.

33. Serbin SP, Singh A, McNeil BE, Kingdon CC, Townsend PA. Spectroscopic determination of leaf morphological and biochemical traits for northern temperate and boreal tree species. Ecological Applications. 2014;24(7):1651–69.

34. Serbin SP, Dillaway DN, Kruger EL, Townsend PA. Leaf optical properties reflect variation in photosynthetic metabolism and its sensitivity to temperature. Journal of Experimental Botany. 2012;63(1):489–502.

35. Kothari S, Beauchamp-Rioux R, Blanchard F, Crofts AL, Girard A, Guilbeault-Mayers X, et al. Predicting leaf traits across functional groups using reflectance spectroscopy. New Phytologist. 2023;238(2):549–66.

36. Burnett AC, Anderson J, Davidson KJ, Ely KS, Lamour J, Li Q, et al. A best-practice guide to predicting plant traits from leaf-level hyperspectral data using partial least squares regression. Journal of Experimental Botany. 2021;72(18):6175–89.

37. Dechant B, Cuntz M, Vohland M, Schulz E, Doktor D. Estimation of photosynthesis traits from leaf reflectance spectra: Correlation to nitrogen content as the dominant mechanism. Remote Sensing of Environment. 2017;196:279–92.

38. Lamour J, Davidson KJ, Ely KS, Anderson JA, Rogers A, Wu J, et al. Rapid estimation of photosynthetic leaf traits of tropical plants in diverse environmental conditions using reflectance spectroscopy. Plos one. 2021;16(10):e0258791.

39. Doughty CE, Santos-Andrade P, Goldsmith G, Blonder B, Shenkin A, Bentley L, et al. Can leaf spectroscopy predict leaf and forest traits along a Peruvian tropical forest elevation gradient? Journal of Geophysical Research: Biogeosciences. 2017;122(11):2952–65.

40. Ely KS, Burnett AC, Lieberman-Cribbin W, Serbin SP, Rogers A. Spectroscopy can predict key leaf traits associated with source–sink balance and carbon–nitrogen status. Journal of experimental botany. 2019;70(6):1789–99.

41. Vasseur F, Cornet D, Beurier G, Messier J, Rouan L, Bresson J, et al. A perspective on plant phenomics: coupling deep learning and near-infrared spectroscopy. Frontiers in Plant Science. 2022;13:836488.

42. Silva-Perez V, Molero G, Serbin SP, Condon AG, Reynolds MP, Furbank RT, et al. Hyperspectral reflectance as a tool to measure biochemical and physiological traits in wheat. Journal of Experimental Botany. 2018;69(3):483–96.

43. Yendrek CR, Tomaz T, Montes CM, Cao Y, Morse AM, Brown PJ, et al. High-throughput phenotyping of maize leaf physiological and biochemical traits using hyperspectral reflectance. Plant physiology. 2017;173(1):614–26.

44. Heckmann D, Schlüter U, Weber AP. Machine learning techniques for predicting crop photosynthetic capacity from leaf reflectance spectra. Molecular plant. 2017;10(6):878–90.

45. Burnett AC, Serbin SP, Rogers A. Source: sink imbalance detected with leaf-and canopy-level spectroscopy in a field-grown crop. Plant, Cell & Environment. 2021;44(8):2466–79.

46. Meacham-Hensold K, Montes CM, Wu J, Guan K, Fu P, Ainsworth EA, et al. High-throughput field phenotyping using hyperspectral reflectance and partial least squares regression (PLSR) reveals genetic modifications to photosynthetic capacity. Remote Sensing of Environment. 2019;231:111176.

47. Fu P, Meacham-Hensold K, Guan K, Wu J, Bernacchi C. Estimating photosynthetic traits from reflectance spectra: a synthesis of spectral indices, numerical inversion, and partial least square regression. Plant, Cell & Environment. 2020;43(5):1241–58.

48. Macklin SC, Mariani RO, Young EN, Kish R, Cathline KA, Robertson G, et al. Intraspecific leaf trait variation across and within five common wine grape varieties. Plants. 2022;11(20):2792.

49. Martin AR, Mariani RO, Cathline KA, Duncan M, Paroshy NJ, Robertson G. Soil compaction drives an intra-genotype leaf economics spectrum in wine grapes. Agriculture. 2022;12(10):1675.

50. Perez-Harguindeguy N, Diaz S, Garnier E, Lavorel S, Poorter H, Jaureguiberry P, et al. Corrigendum to: New handbook for standardised measurement of plant functional traits worldwide. Australian Journal of botany. 2016;64(8):715–6.

51. Gregory LM, Roze LV, Walker BJ. Increased activity of core photorespiratory enzymes and CO2 transfer conductances are associated with higher and more optimal photosynthetic rates under elevated temperatures in the extremophile *Rhazya stricta*. Plant, Cell & Environment. 2023;46(12):3704–20.

52. McClain AM, Sharkey TD. Rapid CO_2_ changes cause oscillations in photosynthesis that implicate PSI acceptor-side limitations. Journal of Experimental Botany. 2023;74(10):3163–73.

53. Stinziano JR, McDermitt DK, Lynch DJ, Saathoff AJ, Morgan PB, Hanson DT. The rapid A/C i response. New Phytologist. 2019;221(2):625–7.

54. Duursma RA. Plantecophys-an R package for analysing and modelling leaf gas exchange data. PloS one. 2015;10(11):e0143346.

55. Delignette-Muller ML, Dutang C. fitdistrplus: An R package for fitting distributions. Journal of statistical software. 2015;64:1–34.

56. Lamour J, Serbin S. spectratrait: A simple add-on package to aid in the fitting of leaf- level spectra-trait PLSR models. In: 1.2.1 Rpv, editor. 2023.

57. Liland K, Mevik B, Wehrens R. pls: Partial Least Squares and Principal Component Regression. In: 2.8-1 Rpv, editor. 2022.

58. Wolkovich E, García de Cortázar-Atauri I, Morales-Castilla I, Nicholas K, Lacombe T. From Pinot to Xinomavro in the world’s future wine-growing regions. Nature Climate Change. 2018;8(1):29–37.

